# A high-quality *de novo* genome assembly of Asian Crested Ibis (*Nipponia Nippon*) using long-read and Hi-C data

**DOI:** 10.1101/2024.04.29.591545

**Authors:** Youngseok Yu, Sung-jin Kim, Changhan Yoon, Jihun Bhak, Changjae Kim, Hyebin Park, Younghui Kang, Yeonkyung Kim, Yu-jin Lee, Seung-yeon Kang, Yong-un Shin, Jong Bhak, Sungwon Jeon

**Author notes:** Correspondence to (Jong B.) and (S. J.). These authors contributed equally to this work (Youngseok Yu, Sung-jin Kim).

## Abstract

We present TtaoRef1, the highest-quality *de novo* genome assembly of Asian Crested Ibis (*Nipponia Nippon*) to date consisting of 134 scaffolds with a length of 1.25 Gb and N50 of 101,183,595 bp. This assembly was generated through the utilization of long-read sequencing and Hi-C data. The assessment of assembly quality, conducted via Benchmarking Universal Single-Copy Orthologs (BUSCO), revealed the presence of 96.8% of completely predicted single-copy genes. TtaoRef1 had 18 times longer N50 value than the previous assembly (ASM70822v1), Furthermore, we conducted the annotation of 24,681 protein-coding genes within the newly assembled genome sequences.

## Data description

### Background and context

The Asian Crested Ibis (*Nipponia Nippon;* Ttaogi, Korean pronunciation) is one of the most endangered bird species in the world [1][2]. It once inhabited vast regions of Northeast Asia, spanning China, Russia, Korea, and Japan until the 1860s. However, due to extensive over-hunting and habitat loss, crested ibis populations experienced a severe decline during the late 19th and early 20th centuries, nearly leading to their complete extinction in the wild. In 1981, seven crested ibises comprising two breeding couples (four adults, with one couple successfully raising three chicks) were discovered in China (Yangxian County, Shanxi). To safeguard these remaining birds, China promptly took comprehensive conservation efforts for both wild and captive populations. Subsequently, two breeding pairs became pivotal in the recovery of the crested ibis population, representing a modern-day ‘Adam and Eve’ for the species. Presently, successful recovery efforts have reintroduced the species in Korea [2]. The last wild Asian Crested Ibis in Korea was extinct since 1978. Over 360 crested ibises have been rehabilitated in the restoration center in Changnyeong, Korea. However, the recovery has been limited to a small number of parent birds, potentially resulting in low biological diversity. This raises the necessity for accurate measurement of genomic diversity within the recovered population. A meticulous assessment of genomic diversity holds crucial implications for the ongoing recovery and restoration of this species, both in Korea and globally. In 2014, the Avian Genome Consortium reported a genome assembly of Asian Crested Ibis, revealing a total sequence length of 1.22Gb [3]. However, the reported genome assembly was based on short-read sequencing data, resulting in low contig N50 values, which clearly delineates its limitations and underscores the need for methodological advancements.

To construct a higher quality Asian Crested Ibis reference genome, we conducted comprehensive whole-genome sequencing (WGS) using both long-read and short-read technologies from a single individual of Asian Crested ibis from Upo Wetland Ecological Park, Changnyeong, Korea. The long-read WGS data, generated through the PromethION platform of Oxford Nanopore Technologies (ONT), was used to construct high-quality primary genome assemblies. To enhance accuracy of the bases, the assembled contigs were further corrected by the short reads sequenced by the MGI-T7 platform of MGI. Additionally, Hi-C data were generated to build scaffolds from the contigs. This improved genome assembly not only achieves better contiguity than ASM70822’s through the utilization of long-read sequencing data for backbone contigs but also enhances the accuracy of predicted gene sequences and structures. These advancements contribute to a better understanding of the genomics of the Asian Crested Ibis and facilitate population-scale genomic studies of this species.

## Methods

### Sample preparation

The blood samples were collected from an Asian Crested Ibis in Upo Wetlands, Changnyeong-gun, Republic of Korea. Genomic DNA was extracted from the Asian Crested Ibis blood samples using the DNeasy Blood & Tissue Kit from QIAGEN according to the manufacturer’s protocol. The quality and concentration of the extracted DNA were evaluated using Epoch Microplate Spectrophotometer and Qubit 4 Fluorometer (Thermo Scientific). Total RNA was extracted using a PAXgene blood RNA kit from Qiagen (Qiagen, USA), according to the manufacturer’s recommendations. RNA quality was assessed by running 1 μl on the 4200 TapeStation system (Agilent, Santa Clara, CA, USA) to ensure RIN and rRNA ratio.

### Short-read and Long-read sequencing

Library construction and whole genome sequencing were conducted by the DNBSEQ-T7 platform of MGI Tech Co., Ltd. (MGI) using the DNBseq™ short-read 150bp paired-end sequencing. Sequencing libraries for long reads were prepared using the standard ligation sequencing kit (SQK-LSK110) (Oxford Nanopore Technologies, UK) following the manufacturer’s instructions. The products were quantified using the 4200 TapeStation system (Agilent, Santa Clara, CA, USA). For long-read sequencing, the raw signal data were generated on the PromethION P48 platform (Oxford Nanopore Technologies, UK). Base-calling from the raw signal data was carried out using MinKNOW v21.11.7 with the Flip-Flop hac model (Oxford Nanopore Technologies, UK).

### Hi-C sequencing data generation

Hi-C chromosome conformation capture data were generated employing the Arima High Coverage HiC kit (A160105 v01, San Diego, CA, USA), with the chromatin digestion carried out using four restriction enzymes (DpnII, HinfI, DdeI, and TruI). For the preparation of Asian Crested Ibis samples for Hi-C analysis, peripheral blood mononuclear cells were extracted from the blood, cross-linked following the manufacturer’s instructions. Subsequently, 0.5 million cross-linked cells were utilized as input in the Hi-C protocol. In brief, chromatin from cross-linked cells was solubilized and subjected to digestion using the four specified restriction enzymes. The spatially proximal digested ends were labeled with a biotinylated nucleotide and subsequently ligated to capture both the sequence and structural information of the genome. Ligation products were purified, fragmented, and size-selected using AMpure XP Beads. Biotinylated fragments were then enriched utilizing Enrichment beads, and Illumina-compatible sequencing libraries were constructed on end repair, dA-tailing, and adapter ligation using a modified workflow derived from the SureSelect XT HS2 Library Preparation Kit (5500-0147, Agilent Technologies, Inc.). The bead-bound library was then amplified, and the resulting amplicons were purified using AMpure XP beads. Validation of the final libraries was performed through qPCR and Bioanalyzer. Subsequently, the libraries were sequenced on the Illumina NovaSeq 6000 platform with 150bp paired-end format at ∼100× coverage following the manufacturer’s protocols.

### RNA-sequencing

We employed 100 ng of total RNA extracted from the Asian Crested Ibis to prepare the sequencing library by using the TruSeq RNA sample preparation kit (Illumina, CA, USA). The quality assessment of the cDNA library was conducted with the 4200 TapeStation system (Agilent, Santa Clara, CA, USA). The library was quantified with the KAPA library quantification kit (Kapa Biosystems, MA, USA) following the manufacturer’s library quantification protocol. After the denatured templates underwent cluster amplification, sequencing was executed in a paired-end (2×100bp) format on the Illumina NovaSeq6000 S4 platform.

### *de novo* assembly of the Ibis genome

We performed quality control and filtering of the raw sequencing reads by removing ten bases at the 5’ end and low-quality base pairs in short-reads from MGI-T7 and Hi-C sequencing reads using Cutadapt 3.7 default options [4]. To estimate genome size, we conducted K-mer analysis using different Ks (from 15-mer to 36-mer) using the Jellyfish program [5]. The filtered reads were then used to construct the *de novo* assembly. To achieve the highest quality Ibis genome assembly utilizing both nanopore long reads and short reads, we employed five distinct *de novo* assembly programs (wtdbg2, canu v.2.2, flye v.2.9.2, NECAT and NextDenovo v.2.5.0) and two polishing programs (pilon and NextPolish v.1.4.1) [6-11]. Then, we scaffolded the assembly (NextDenovo v.2.5.0+ NextPolish v.1.4.1) using Hi-C data by SALSA tools [12], which showed the optimal contiguity. The completeness of the *de novo* assembly of the Ibis genome was evaluated using the Benchmarking sets of Universal Single-Copy Orthologs (BUSCO) v.5.4.5 program. [13]

### Assembly sequence comparison

Whole-genome alignment was conducted to investigate the sequence similarity between TtaoRef1 and ASM70822. We used Nucmer (–maxmatch -b 500 -c 500 -l 100) and mummerplot (mummerplot --png --filter -fat --large --layout -R $Ref -Q $Query -l) from Mummer4 (version 4.0.0rc1) to align the existing assembly to the new one and to generate a dotplot [14]. The genomic differences were presented by passing the nucmer delta file to dnadiff, a tool from Mummer4 (with a default option). Numbers of SNPs and INDELs were calculated as gSNPs and gINDELs from dnadiff, which was supported by at least 20 exact matches at both ends of each variant site. To visually compare the contiguity between assemblies, the delta file was filtered using delta-filter (-1 -i 90 -l 100), a tool from Mummer4, to remove small and lower quality alignments. The filtered delta was then visualized using circos (v.0.69-9). [15]

### Genome annotation

To generate supporting evidence of the gene structure, we first obtained quality controlled RNA sequence and filtered out ten bases at the end, low-quality bases pairs and adapter sequences in raw RNA reads from MGI-T7 sequencing using the Fastp program default options [16]. Then, we mapped transcripts using STAR v.2.7.1a [17] against the assembled Ibis genome. We identified and masked repeats using RepeatMasker v4.0.5 [18]. Finally, We applied the BRAKER2 tool [19] with ‘--useexisting –specific=chicken –softmasking’ option with protein coding sequence from ASM70822y [3] and the RNA-seq mapping information to predict protein-coding genes. The completeness of the predicted transcript sequences was evaluated using the BUSCO v.5.4.5 [13].

## Data validation

### Whole-genome sequencing and K-mer analysis

We generated 123.34 Gb of long-read sequences from a single female Asian crested Ibis individual with N50 of 27.35 Kb by the PromethION platform of Oxford Nanopore Technologies (ONT). Concurrently, a total of 204.01Gb short-read WGS data of the same individual were generated by the MGI-T7 platform of MGI. Following quality control and trimming, the filtered short-read data had 190.08 Gb. Hi-C data were also generated for the same individual, totaling 167.53 Gb. K-mer analysis showed that the genome size of Asian Crested Ibis was estimated to be 1.11 Gb (Figure 1), aligning closely with the ASM70822[3].

**Figure 1.**
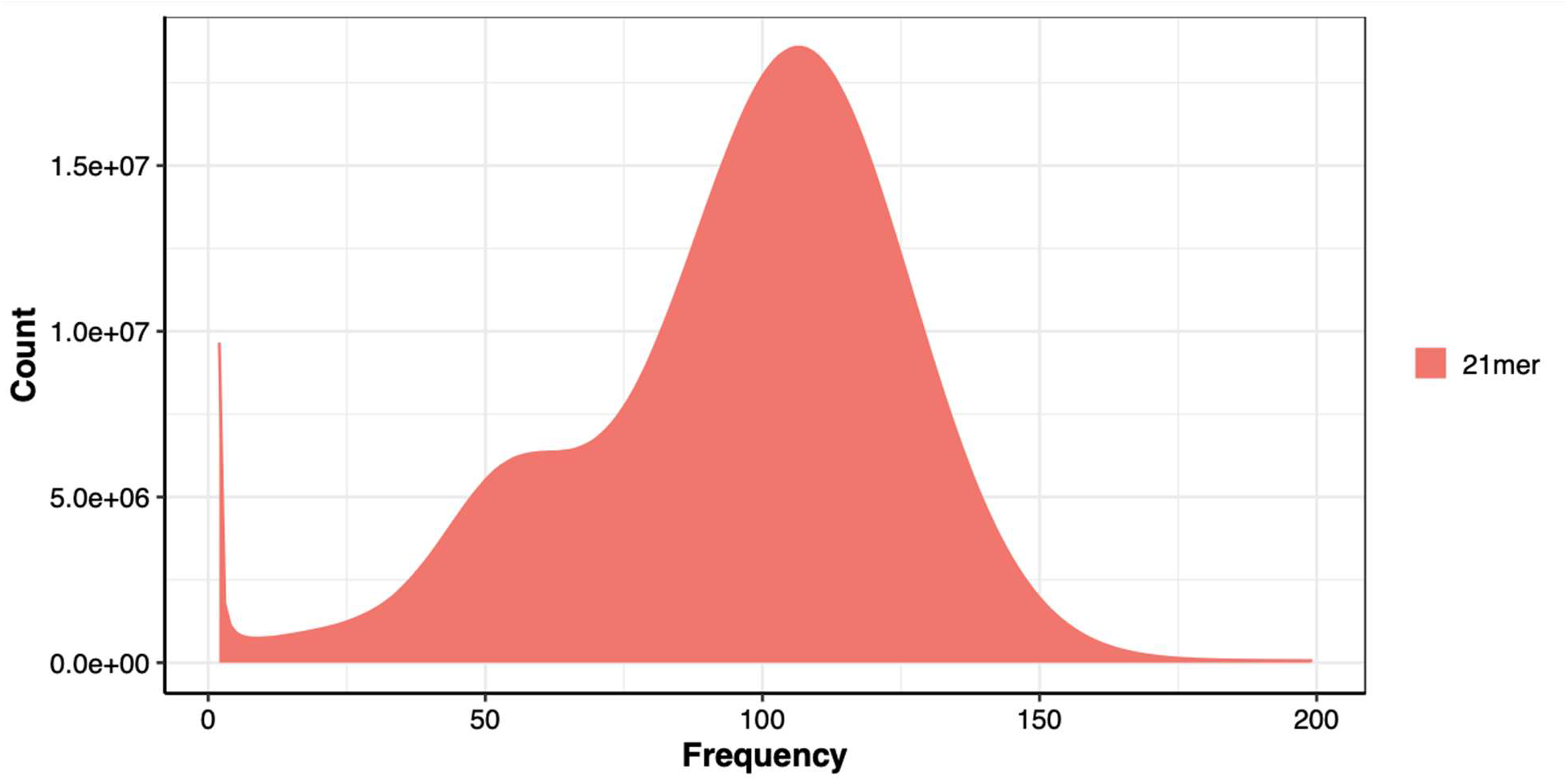
K-mer analysis results of the Asian crested Ibis using K=21.

## Construction of *de novo* genome assembly

A total of 1,248,612,181 bases of *de novo* genome assembly of Asian Crested Ibis was generated using the Nextdenovo tool (v.2.5.0). We initially constructed five *de novo* assemblies with five distinct combinations of tools and options (Table 1). The NextDenovo+NextPolish approach demonstrated the highest assembly quality regarding scaffold N50 value (101,183,595 bp) and the minimal number of assembled contigs (177). We further assessed the quality of the assemblies using BUSCO, and NextDenovo + NextPolish had 96.8% of complete single-copy genes from BUSCO, indicating the high quality of the *de novo* assembly (Table 2). Subsequently, contigs were scaffolded into 134 scaffolds using Hi-C data by SALSA. The final *de novo* genome assembly comprised 1,241,683,096 bases (Table 1)

**Table 1.** Quality metrics of 5 assemblies of *Nipponia Nippon*.

|  | wtdbg2 +<br>pilon | canu +<br>pilon | flye + pilon | NECAT +<br>pilon | NextDenov<br>o +<br>NextPolish | NextDenov<br>o +<br>NextPolish<br>+ Salsa |
| --- | --- | --- | --- | --- | --- | --- |
| Number<br>of<br>Contigs | 1,549 | 1,249 | 1,021 | 902 | 177 | 134 |
| Assemb<br>ly<br>length<br>(bp) | 1,201,657,619 | 1,324,695,631 | 1,242,037,173 | 1,321,730,832 | 1,248,612,181 | 1,241,683,096 |
| Max of<br>contig | 83,572,929 | 37,933,964 | 88,713,641 | 114,528,015 | 131,586,178 | 167,974,882 |
| N50<br>(bp) | 43,626,617 | 14,265,810 | 28,282,931 | 42,821,202 | 56,735,934 | 101,183,595 |
| N60<br>(bp) | 28,102,841 | 11,897,818 | 18,290,105 | 26,483,896 | 30,429,921 | 54,427,359 |
| N70<br>(bp) | 21,497,698 | 7,211,952 | 13,041,921 | 17,156,712 | 24,124,264 | 49,745,395 |
| N80<br>(bp) | 12,307,138 | 4,476,561 | 8,854,853 | 11,988,719 | 13,263,679 | 17,519,629 |
| Mean<br>length<br>(bp) | 775,764 | 1,060,605 | 1,216,491 | 1,465,334 | 7,054,306 | 9,318,202 |
| Median<br>length<br>(bp) | 15,733 | 84,234 | 11,753 | 61,354.50 | 353,914 | 377,558 |

**Table 2.**
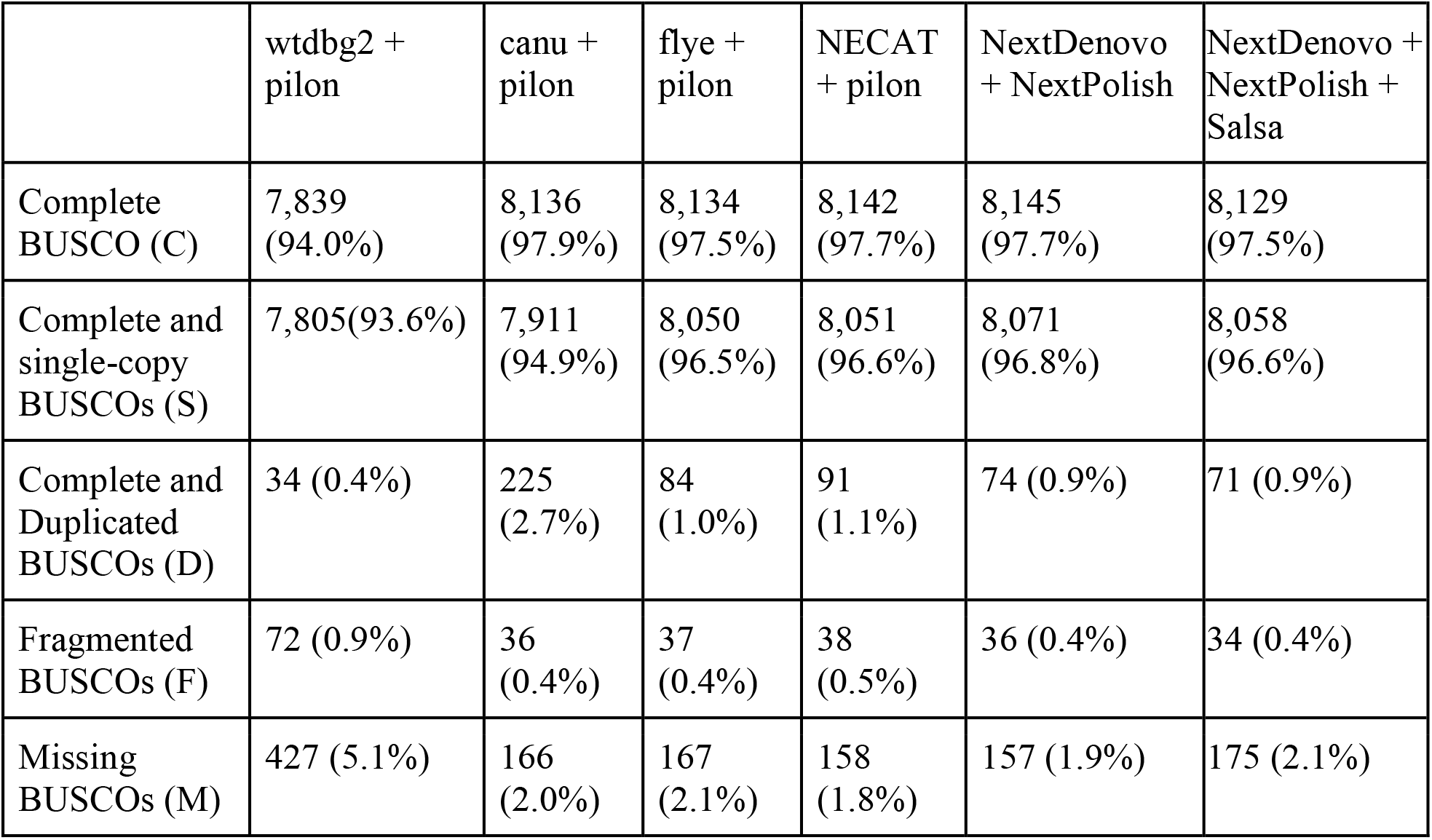
BUSCO result metrics in aves_odb10 database of 5 assemblies of *Nipponia Nippon*.

|  | wtdbg2 +<br>pilon | canu +<br>pilon | flye +<br>pilon | NECAT<br>+ pilon | NextDenovo<br>+ NextPolish | NextDenovo +<br>NextPolish +<br>Salsa |
| --- | --- | --- | --- | --- | --- | --- |
| Complete<br>BUSCO (C) | 7,839<br>(94.0%) | 8,136<br>(97.9%) | 8,134<br>(97.5%) | 8,142<br>(97.7%) | 8,145<br>(97.7%) | 8,129<br>(97.5%) |
| Complete and<br>single-copy<br>BUSCOs (S) | 7,805(93.6%) | 7,911<br>(94.9%) | 8,050<br>(96.5%) | 8,051<br>(96.6%) | 8,071<br>(96.8%) | 8,058<br>(96.6%) |
| Complete and<br>Duplicated<br>BUSCOs (D) | 34 (0.4%) | 225<br>(2.7%) | 84<br>(1.0%) | 91<br>(1.1%) | 74 (0.9%) | 71 (0.9%) |
| Fragmented<br>BUSCOs (F) | 72 (0.9%) | 36<br>(0.4%) | 37<br>(0.4%) | 38<br>(0.5%) | 36 (0.4%) | 34 (0.4%) |
| Missing<br>BUSCOs (M) | 427 (5.1%) | 166<br>(2.0%) | 167<br>(2.1%) | 158<br>(1.8%) | 157 (1.9%) | 175 (2.1%) |

### Comparison to the previous genome assembly

TtaoRef1 demonstrated a significantly improved scaffold N50 value, being 19 times longer at 101.18 Mb, in comparison to ASM70822 with an N50: of 5.21 Mb, while maintaining a comparable genome size. TtaoRef1 exhibits superior contiguity, as evident in key metrics such as the N50 length, the longest scaffold length, and the number of scaffolds (Table 3). Notably, we identified 40 Mb of novel sequences that were not present in ASM70822. Moreover, the comparative analysis revealed 46,710 links, signifying a homologous relationship between both assemblies, which were derived from the scaffolds larger than 10Kb. The Circos plot visually depicts the better contiguity of TtaoRef1 compared to ASM70822 while effectively covering the majority of the fragmented scaffolds from ASM70822 (Figure 2).

**Table 3.** Comparison of the assemblies. SNPs: Single Nucleotide Polymorphism counts; gSNPs: Single Nucleotide Polymorphisms bounded by 20 exact, base-pair matches on both sides; Indels: Single Nucleotide Insertions/Deletions; gIndels: Single Nucleotide Insertions/Deletions bounded by 20 exact, base-pair matches on both sides; Inversions: Number of breaks in the alignment where adjacent 1-to-1 alignment blocks are inverted with respect to one another; Relocation: Number of breaks in the alignment where adjacent 1-to-1 alignment blocks are in the same sequence, but not consistently ordered; Translocation: Number of breaks in the alignment where adjacent 1-to-1 alignment blocks are in different sequences.

|  | **TtaoRef1 (This study)** | **ASM70822 (Previous study)** |
| --- | --- | --- |
| Total length (bp) | 1,241,683,096 | 1,223,863,029 |
| Scaffold N50 length (bp) | 101,183,595 | 5,211,696 |
| Contig N50 length (bp) | 56,735,934 | 29,116 |
| Max scaffold length (bp) | 167,974,882 | 27,015,053 |
| Number of scaffolds | 134 | 59,555 |
| Covered bases by alignment (bp) | 1,175,184,310 (94.12%) | 1,170,505,885 (95.64%) |
| GC content | 0.4275 | 0.4183 |
| Number of SNPs | 869,066 | 869,066 |
| Number of gSNPs | 455,918 | 455,918 |
| Number of Indels | 4,572,325 | 4,572,325 |
| Number of gIndels | 47,743 | 47,743 |
| Number of Inversions | 139 | 348 |
| Number of Relocations | 1,018 | 1,055 |
| Number of Translocations | 6,120 | 265 |

**Figure 2.**
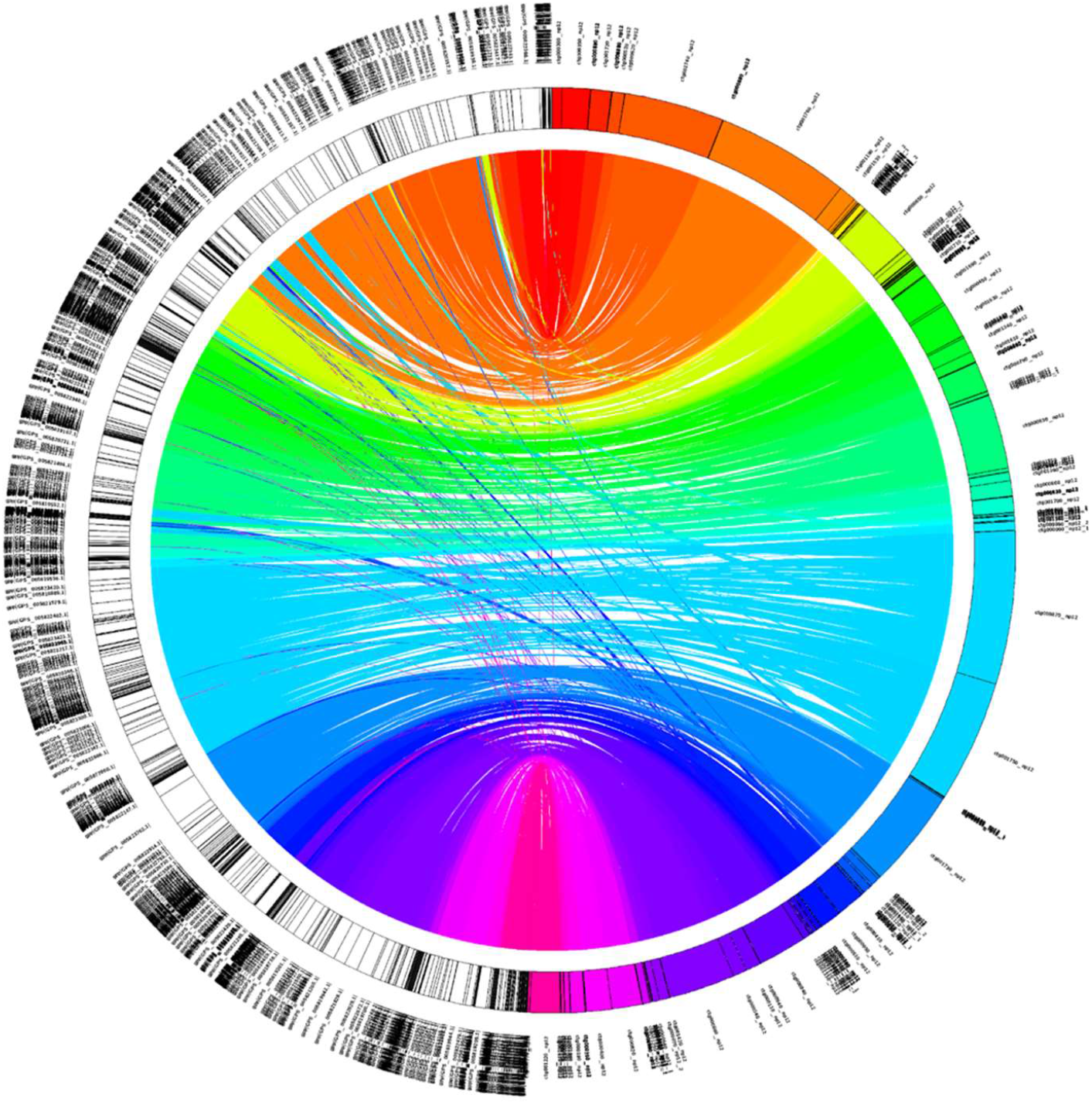
Circos plot of TtaoRef1 in comparison to ASM70822. The right half-circle indicates TtaoRef1

### Genome annotation

We predicted a total of 24,681 genes and 28,268 transcripts from the newly assembled Asian Crested Ibis genome using the BRAKER2 tool [19] and RNA-seq data. The N50 length of the transcripts was determined to be 2,148 bp (Table 4, Figure 3). To evaluate the quality of the predicted genes, we employed BUSCO analysis, showing 88.2% of complete BUSCO (single-copy BUSCOs 73.8% and duplicated BUSCOs 14.4%), indicating the high quality of the genome annotation results.

**Table 4.** Statistics of the predicted transcripts.

|  | Transcript |
| --- | --- |
| Longest of transcript length (bp) | 64,614 |
| Shortest of transcript length (bp) | 12 |
| Median of transcript length (bp) | 999 |
| mean of transcript length (bp) | 1,462 |
| Transcript N50 length (bp) | 2,148 |
| Transcript N60 length (bp) | 1,719 |
| Transcript N70 length (bp) | 1,359 |
| Transcript N80 length (bp) | 1,032 |
| Transcript N90 length (bp) | 687 |

**Figure 3.**
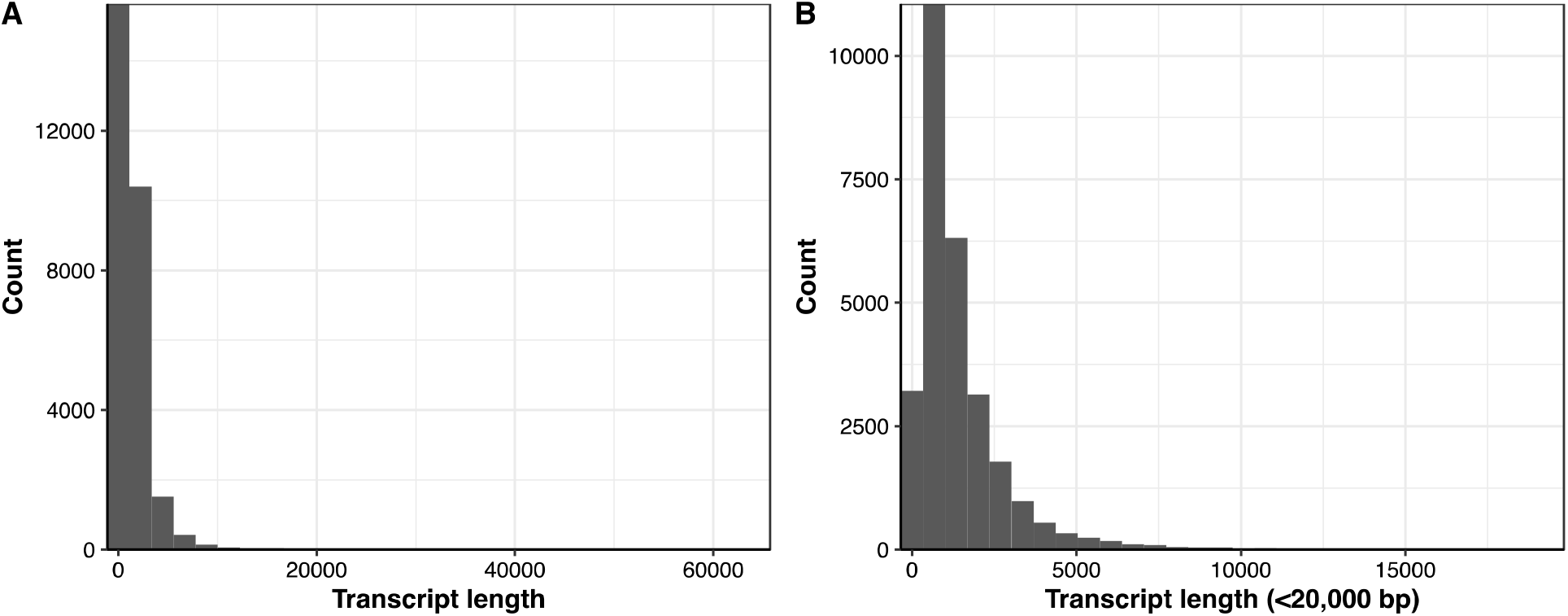
Length distribution of (A) all predicted transcripts and (B) transcripts (< 20,000 bp)

## Conclusion

We generated an improved version of the *de novo* genome assembly for the Asian Crested Ibis, incorporating ONT long-read and short-read whole-genome sequencing data. Along with Hi-C reads, the assembly had a genome size of 1.24Gb and consisted of 134 scaffolds. Notably, TtaoRef1 showed 19 times longer scaffold N50 value than ASM70822 by utilizing long-read sequencing data, indicating a robust sequence contiguity. The Asian Crested Ibis reference genome constructed in this study provides a robust foundation for investigating the population genomic diversity of this species and devising more effective strategies for its recovery.

## Supporting information

Pictures of Asian crested ibis in Upo wet-land area, Changnyeong, Gyeongsangnam-do, Republic of Korea. The pictures were provided by Changnyeong count

## Data Availability

The genome assembly was also deposited at NCBI under project accession number PRJNA1049517 and sample number SAMN38701424.

## Authors’ contributions

Sung-jin. K., Y.L., Seung-yeon. K. and Y. S. provided the blood samples. S. J., Y.Y., C.Y., and Jihun B. conducted bioinformatics analyses. C.K., H.P., Y. Kang, and Y. Kim conducted wet-lab experiments. S. J., Y.Y., C.Y., Jihun B., and C.K., Jong B. wrote and revised the manuscript. Jong B. supervised the study.

---

**Figure S1.**
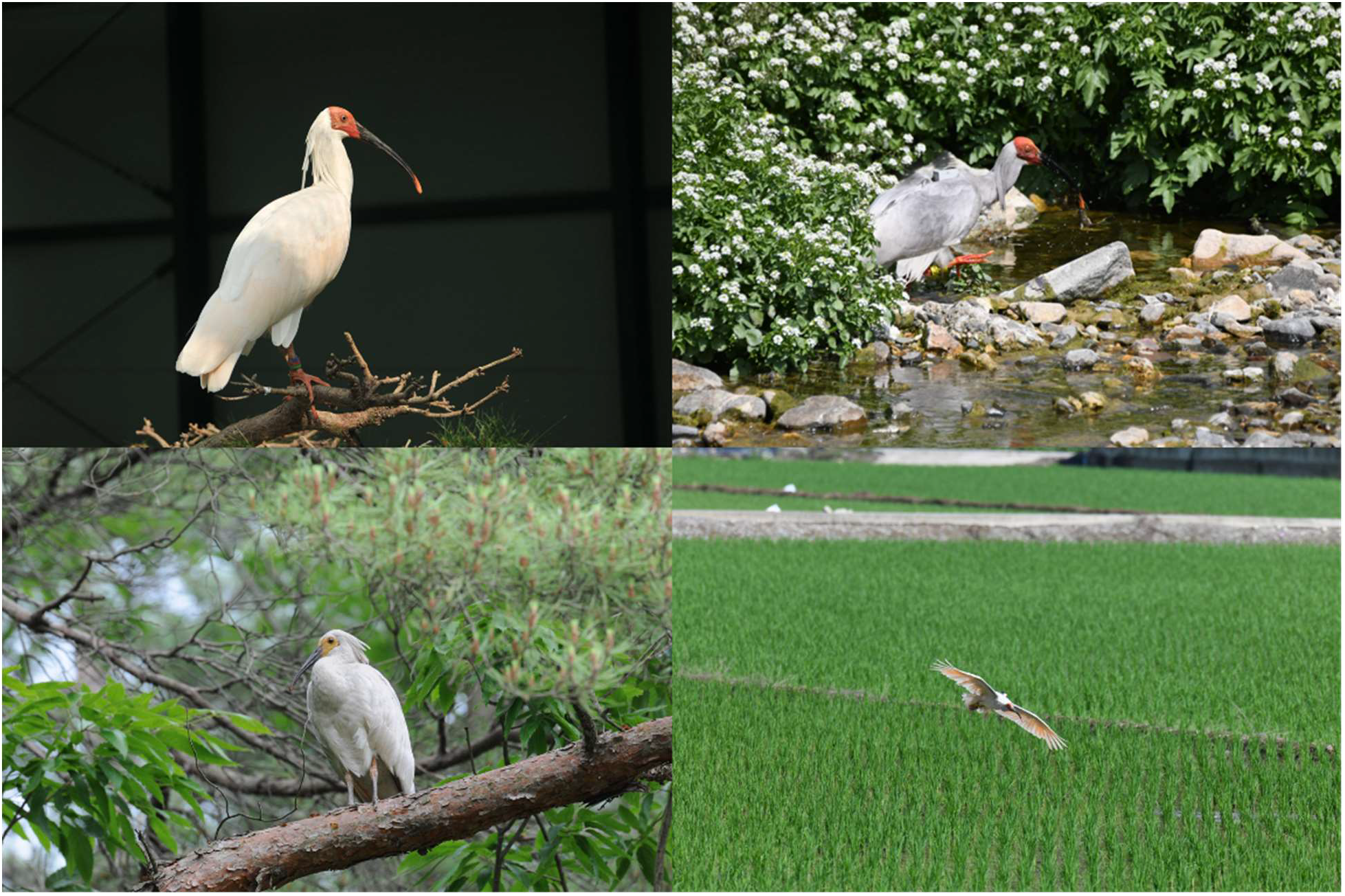
Pictures of Asian crested ibis in Upo wet-land area, Changnyeong, Gyeongsangnam-do, Republic of Korea. The pictures were provided by Changnyeong county.

