## Supplementary material for "A high-quality *de novo* genome assembly of Asian Crested Ibis (*Nipponia Nippon*) using long-read and Hi-C data": Pictures of Asian crested ibis in Upo wet-land area, Changnyeong, Gyeongsangnam-do, Republic of Korea. The pictures were provided by Changnyeong count

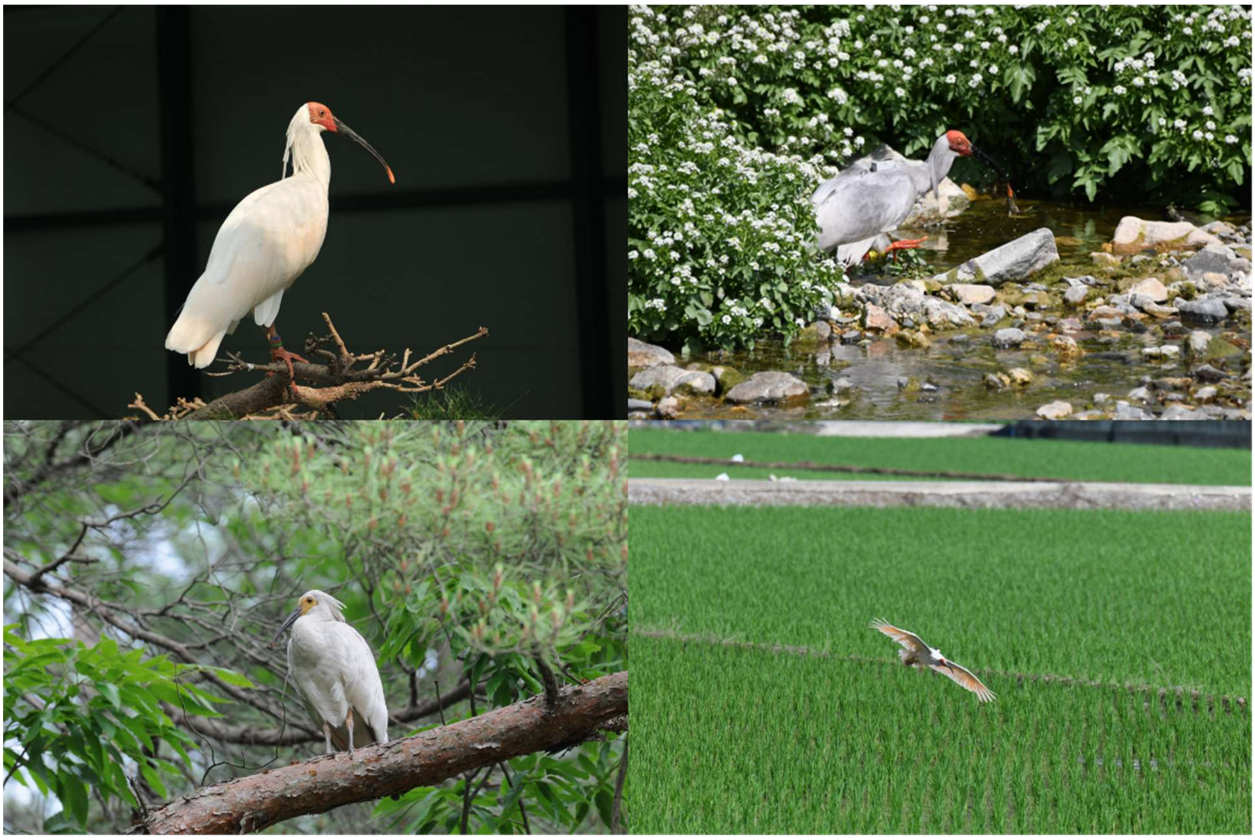

**Figure S1. Pictures of Asian crested ibis in Upo wet-land area, Changnyeong, Gyeongsangnam-do, Republic of Korea.** The pictures were provided by Changnyeong county.
